# Variable sparsity non-negative matrix and tensor factorization algorithms and their applications in cancer genomics

**DOI:** 10.64898/2026.10.19.760022

**Authors:** Timothy Pham, Jiaqian Zhong, Joseph Hwang, Ronit Agarwala, Kun Zhang, Prashant Mali, William Kim, Pablo Tamayo

## Abstract

The field of oncology has experienced significant advancements in novel therapies and molecular diagnostics to personalize initial cancer treatments. However, cancer’s propensity for evolution and resistance to therapies remains a formidable challenge due to its dynamic nature. Cancer’s complex and adaptive behavior surpasses the capabilities of simple models, complicating our comprehension at every level, from basic sciences to clinical practices, highlighting the urgent need for innovative approaches to improve the modeling and classification of cancer. In this context, matrix factorization algorithms have been used to generate transcriptional signatures that summarize the most salient patterns of behaviors in cancer datasets. For example, the standard non-negative matrix factorization algorithm (*NMF*) generates a dimensionality-reduced representation that encodes a significant amount of data relationships as a mixture of positive components and provides a working summary of the input data. The resulting *NMF* factor matrices, *W* and *H*, display some degree of sparseness; however, this sparseness is often a byproduct rather than an explicit objective of the *NMF* algorithm, and the user has limited control over the degree of sparseness of the representation and the corresponding balance between local versus global structures in the *NMF* results. Some sparse versions of *NMF* have been introduced but still do not allow a fine-grained control on the sparsity constraints. We present here a novel *variable-sparsity Non-negative Matrix Factorization algorithm* (*vsNMF*), where instead of imposing sparsity as a regularization constraint on the *NMF* objective function, sparsity is explicitly enforced by a projective step performed for each column of *W* and row of *H* after a standard *NMF* update. In this way, the algorithm takes as input the input matrix, plus two one-dimensional vectors, that specify the sparsity functions that will be imposed on the *NMF* solution. The more explicit control of sparsity, and the corresponding balance between local vs. global structure, at the level of individual components, allows the generation of *NMF* solutions that can be tuned to operate anywhere between a generative, parts-based encoding of overlapping features, and a partitional, soft-clustering model. This allows greater flexibility to represent biological subtypes, patterns of pathway activation or deregulation, and higher biological interpretability. These learned low-dimensionality representations can also be used to integrate and merge datasets, or to define groups of similar cancer samples representing putative cellular or tumor states. Adjustable/tunable *NMF* representations of biological data can greatly help advance our fundamental understanding of cancer biology, provide more interpretable computational models of cancer, and better functional cancer taxonomies and precision oncology paradigms. We apply the same approach to tensors and introduce a variable-sparsity Non-Negative Tensor Factorization algorithm (*vsNTF*) that can take an input tensor and sparsity functions for each of the tensor dimensions.

## Introduction and Motivation

Over the past decade, the field of oncology has witnessed groundbreaking advances in targeted and immuno-therapies, alongside the integration of molecular diagnostics to tailor initial cancer treatments [1,2]. However, the dynamic nature of cancer, characterized by its ability to evolve and resist therapies often mediated by cancer cell plasticity remains a formidable challenge from a modeling perspective. Cancer exhibits the defining characteristics of an emergent complex system [3–12] and challenges the reliance on mutation status as a proxy for cancer “state,” as it often provides limited insights into the dynamic nature of cancer behavior, contrary to conventional belief. Cancer genomics datasets are incredibly high-dimensional. A single tumor sample might be characterized by the expression of 20,000+ genes, millions of methylation sites, or thousands of mutation types. However, the number of available samples, e.g. cell lines, mouse models or patients, is often much smaller. This leads to a “curse of dimensionality,” that makes it difficult to train machine learning models without overfitting. Overcoming these challenges requires developing a useful low-dimensionality representation or embedding and effectively modeling the complex phenomenology of cancer cells. The cancer modeling strategy our lab has pursued over the last decade [13–16] involves making computational models of the observed cellular states and phenomenological traits of cancer, focusing on the collective and empirical characteristics of these complex systems and making models by dimensionality reduction, clustering, projection and using association metrics to compare, and find relationships between cancer sample profiles. In order to maximize the potential benefit of these models the computational techniques emphasize the model’s biological interpretability.

The use of Non-negative Matrix Factorization (*NMF*) has revolutionized cancer genomics by providing a biologically interpretable method for dimensionality reduction and dataset decomposition. Because *NMF* constrains its outputs to non-negative values, it allows researchers to summarize massive, high-dimensional multi-omics datasets into additive, easily understandable components. Most notably, *NMF* has been the foundational algorithm to decompose mRNA datasets into transcriptional components that summarize and catalog common patterns of gene expression, and these “metagenes” have vastly improved tumor classification [13,17–23]. *NMF* has also been used to discover and map “mutational signatures” by decomposing complex tumor mutation catalogs into distinct patterns linked to specific environmental exposures, such as smoking or UV radiation, and internal cellular defects, such as faulty DNA repair [22,24]. In our own work we have used NMF to generate signatures to define subtypes in Multiple Sclerosis [25], characterize ribosomal protein mRNA groups [26], define DNAm components and distinct subtypes of myelodysplastic syndromes [27], and to generate signatures of copy number alterations [21]. One remarkable property of *NMF* is its ability to produce a *dimensionality-reduced* representation of an input dataset that encodes a significant amount of data relationships using only a few positive components, exposing common patterns, and providing a working summary model of the data [17,28–30] that is much more interpretable than those generated by other traditional dimensionality reduction algorithms such as *PCA* or *ICA*. For example, it is advantageous to generate this type of representation for bulk and single cell mRNA profiles because it simplifies modeling and offers interpretability through its non-negative constraints. We and others have used *NMF* to successfully decompose genomic datasets of different types [13,17–23]. However, despite the successful application of *NMF* in many problems in cancer genomics, we believe it will be beneficial to gain additional control of the structure of the *NMF* solution and have additional flexibility to tailor the *NMF* solution to the requirements of the genomics problem at hand. For instance, one may want to “align” the NMF solution (*W* and *H* matrices) in such a way that it better represents “functional” or “response” cell states, namely those relevant to cancer dynamics or perturbation-induced transitions. In other cases, one may want to make the *NMF* solution to better reflect disease subtypes determined by cell of origin, morphology or tissue types. One could also envision modulating an *NMF* solution to be better as providing an intermediate representation to clustering or predictive models. Finally, one may want to tune the *NMF* solution to be more transparent and biologically interpretable and better match profiles of specific gene sets or signatures that represent well understood biology.

In the past there have been attempts to control the *NMF* decomposition by imposing a global degree of sparseness or smoothness by adding regularization terms or structural constraints to the objective function. For example, *sparse NMF* [31] introduces parameters to explicitly control the sparsity of either the *W* or *H* matrices by adding a regularization penalty to the objective function. Smoothing approaches to *NMF* [32,33] penalize large differences between adjacent elements in the basis or activation vectors by adding a regularization term or a penalty based on the second-order derivative [34]. *Graph-Regularized NMF* (*GNMF*) [35] ensures that if two data points are close to each other in the original high-dimensional space, their representations in the new lower-dimensional space (the columns of *H*) are also close by using a data-based nearest-neighbor Laplacian graph as an additional regularization term. *Orthogonal NMF* [36] emphasizes the orthogonality of the basis vectors or the coefficient vectors to be orthogonal by imposing Orthogonality constraints (e.g., *W*^*T*^*W = I* or *HH*^*T*^ *= I*). All these approaches provide some amount of extra control of the *NMF* solution by adding regularization terms or structural constraints to the objective function. However, modifying the objective function with an additional regularization or penalty comes at a price: the resulting *NMF* solution is then a compromise between the original objective function, favoring minimizing the reproduction error, and the additional constraint, favoring properties such as sparseness or smoothness. As a result, the solution cannot fully satisfy the desired constraint. There is the case when by imposing sparseness constraints, one can influence sparseness but lacking explicit control over it. In addition, if one applies the regularization penalty to only one of the matrices, the *NMF* optimization process may converge on the trivial solution by simply scaling the elements of the matrix toward zero. To prevent this, the algorithms must enforce explicit normalization constraints. This adds another layer of complexity to the optimization process and often requires specialized solvers such as projected gradient descent. Adding complex structural constraints, as in the case of the Graph Laplacian for smoothness, could generate a much more rugged loss landscape making the algorithm significantly more sensitive to its initialization state and potentially more likely to get trapped in suboptimal local minima.

We introduce here a *variable-sparsity Non-negative Matrix Factorization* algorithm (*vsNMF*), that preserves the reproduction error minimization objective function while providing explicit fine-grained control over the sparseness of the *NMF* solution. This allows much greater control of the specific mixture of global versus local structures and the ability to tailor the *NMF* solution for different genomics data use cases. Our *vsNMF* approach is an extension of the original approach of Hoyer [37], where instead of imposing sparsity as a regularization constraint on the *NMF* objective function, sparsity is explicitly enforced by a projective step performed for each column of *W* and row of *H* after a standard *NMF* update. In this way, the algorithm takes as input the input matrix *A*_*nm*_, plus two one-dimensional vectors, *S*^*W*^_*k*_ and *S*^*H*^_*k*_ that specify the sparsity functions that will be imposed on the *NMF* solution. The practical consequence of this fine-grained sparsity control is direct modulation over the *NMF* algorithm’s dual tendency to behave as a parts-based encoder or as a soft clustering method. In *vsNMF*, this tendency is no longer a property of the whole factorization but a per-component setting, as each component can be independently positioned along a continuum between an extreme-ray regime, i.e. a generative, additive parts/encoding, and a centroid-like regime, i.e. partitional, prototype/soft-clustering one [38–40]. This flexibility provides an unprecedented degree of control to make an *NMF* solution better represent several levels or types of biological structure or characteristics in the genomics data. We apply the same approach to tensors and introduce a *variable-sparsity Non-Negative Tensor Factorization* algorithm (*vsNTF*) that can take an input tensor and sparsity functions for each of the tensor dimensions. We will describe the algorithms below and analyze how imposing specific sparseness constraints to the *NMF* decomposition algorithm provides useful solutions for genomics data analysis. In what follows, we will first describe the *vsNMF* and *vsNTF* algorithms in detail and then present some genomics data applications.

### The *variable-sparsity Non-Negative Matrix Factorization* algorithm (*vsNMF*)

The standard *NMF* algorithm is based on the *factorization* of an input or *reference dataset* into a product of two lower-rank matrices [29],

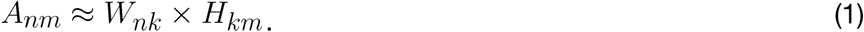

The *W*_*nk*_ matrix contains the positive weights of each *relevant gene* for each *component* or *signature*, and the *H*_*km*_ matrix corresponds to a positive “mixture” of signatures, namely *NMF* components or latent factors, that approximates each reference dataset *A*_*nm*_ column. In equation (1) and for genomics data, *n* is typically the number of *genes, m* is the number of samples, and *k* is the number of *NMF* components, where *k* is much less than *n* or *m*. A numerical solution to this problem typically minimizes the Euclidean error *E(W, H)*, between the original dataset *A*_*nm*_ and its reconstructed version (*W*_*nk*_ *x H*_*km*_),

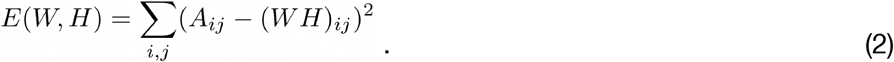

The *W* and *H* matrices are initialized with random values and the decomposition results from applying a sequence of iterative updates (*t* = 0, 1, 2…) in the direction that decreases the Euclidean error [29,41,42],

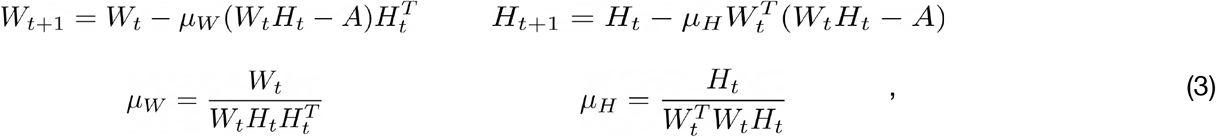

until the algorithm converges to a low error solution. This solution is a local minimum and therefore an approximation to the global solution. Finding the global minimum is an NP-hard problem [43].

The resulting summary matrices *W* and *H* display some degree of sparseness; however, this sparseness is often a byproduct rather than an explicit objective of the algorithm. The user has limited control over the degree of sparseness of the representation and the corresponding balance between local vs. global structure in the *NMF* results. To improve the control of this balance, and in this way select solutions with potentially better representation of biological subtypes and higher biological interpretability, we designed a new *vsNMF* algorithm. The *vsNMF* algorithm generalizes the approach of Hoyer [37], and imposes pre-defined sparsity constraints on the rows of *H* and/or the columns of *W*, where the sparsity *S(x)* for an *H* row (*W* column) vector *x* of *n* elements is measured by how much the ratio between the *L*_*1*_ (manhattan) and *L*_*2*_ (euclidean) norms deviates from its dense limit 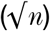 [44,45]. Under this approach, we implemented a modified *NMF* iterative gradient descent algorithm [37], which we refer to as *variable-sparsity NMF* (*vsNMF*), in which the algorithm’s iterations (Fig. 1) are constrained to produce intermediate *W* and *H* matrices by obeying preset values of a sparsity function, *S(x)*,

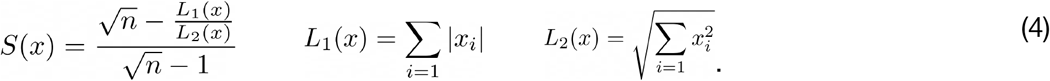

**Figure 1.**
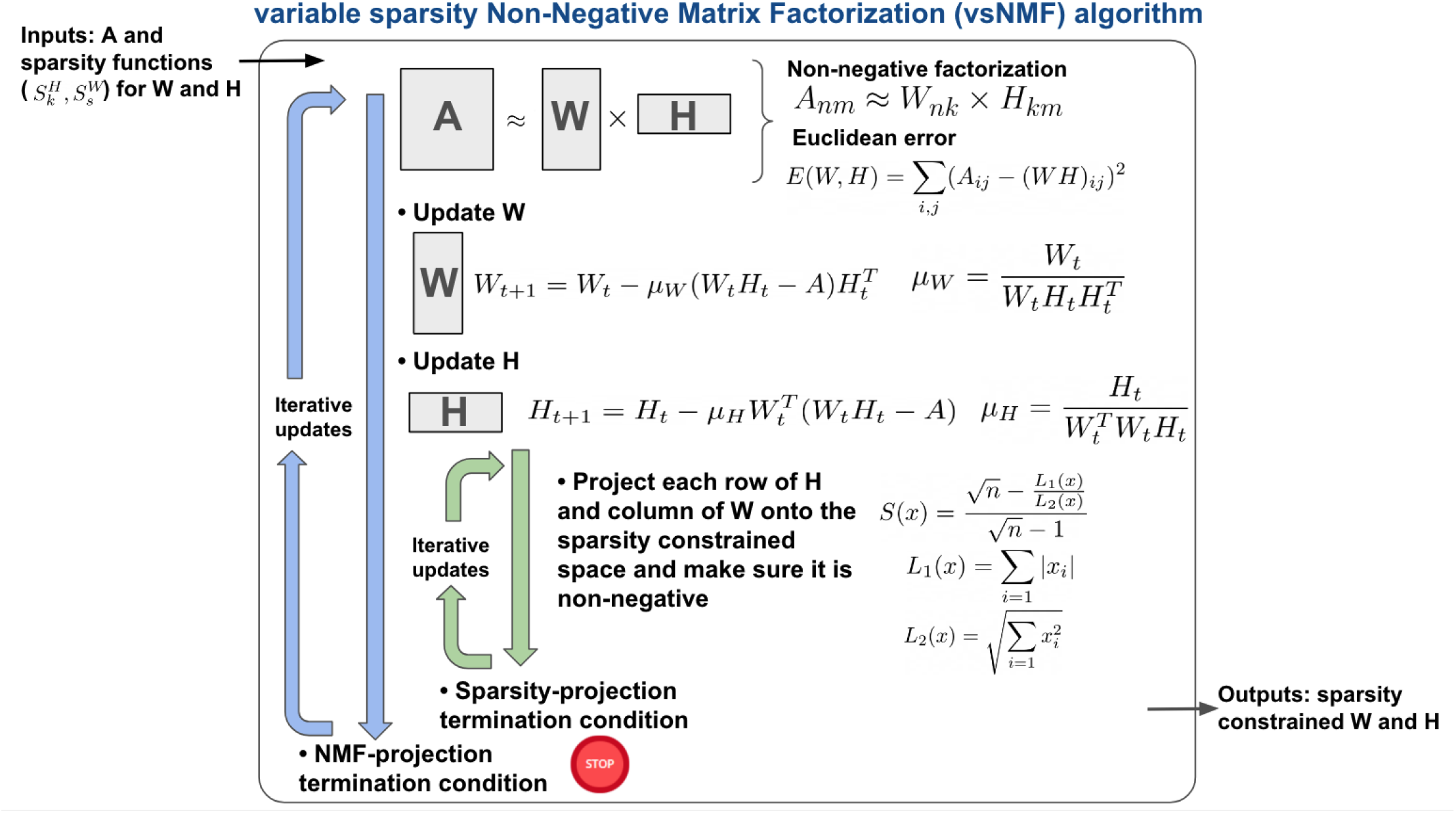
Top-level description of the *vsNMF* decomposition algorithm.

In this manner, independent values of sparsity for each row of *H* or column of *W* can be pre-defined in advance. The preset sparsity values for each row are imposed at each iteration of the algorithm following an update of *H*, in the direction of the negative gradient (eq. 3), with an additional *projection step* to map the results of the *H*-update onto the appropriate non-negative sparsity-constrained space [37]. It is advantageous to define *W* and *H* matrices with different degrees of sparsity by choosing specific choices of the *S*^*H*^_*k*_ and *S*^*W*^_*k*_ functions. This enables, for example, capturing biological patterns at different scales and detecting more fine grained patterns that are less common in standard *NMF* solutions. An interesting choice of sparsity function is a power-law distribution of *H*-matrix sparsities,

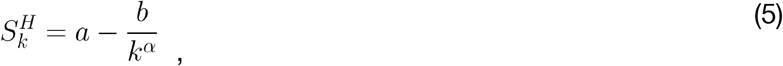

and leaving *W* unconstrained in the *vsNMF* iterations. Using this approach, *a, b* and *α* will be chosen to specify the minimum and maximum sparsity and the specific decay of the power law, respectively. Using these constraints we can produce an *H* matrix structure with more emphasis on localized structure than the original *NMF* algorithm, which produces a roughly uniform distribution of sparsities. This specific sparsity structure also allows us to capture a hierarchical representation of the data where a few *NMF* components represent top global behavior and at the same time the remaining ones are dedicated to representing intermediate and more local behavior of the samples further down the implicit hierarchy of cell activity. This approach increases the capability of *NMF* to capture biologically-relevant patterns at different scales such as those relevant to summarize behavior at the local or global level and define, e.g., cellular or tumor states.

While the application examples below present detailed results of using the *vsNMF* algorithm to decompose genomic data, we will first provide two simple examples demonstrating the algorithm’s output and the effects of modifying the sparsity function across two input datasets. The first is the ORL faces dataset, also known as the AT&T faces dataset, a widely used benchmark in face recognition research created between 1992 and 1994 at the Olivetti Research Laboratory in Cambridge [46]. This dataset contains 400 grayscale images of 40 distinct subjects (10 images per person), with each image having a resolution of 92×112 pixels. The images were captured at different times under varying lighting conditions, with diverse facial expressions, e.g., open or closed eyes, smiling or not smiling), and details, e.g., with or without glasses, against a dark, homogeneous background. Because they were taken by a human photographer, the images also exhibit slight variations in scale and orientation. The second dataset is the CCLE solid tumor cancer cell lines dataset, a subset of mRNA profiles from the Cancer Cell Line Encyclopedia [47,48] that contains 3,205 genes representing pan-cancer oncogenic pathways across 1,086 samples from over 20 different cancer types. This will also serve as the first dataset in the applications section below. As shown in Fig. 2, imposing different sparsity constraints significantly impacts the types of solutions generated by *vsNMF*. For the faces dataset, we apply sparsity constraints to the *W* matrix. Because this matrix represents image components that act as building blocks, e.g., facial features like eyes, ears, noses, and mouths, it is desirable to control the degree of wholeness or locality of these characteristics. Conversely, for the CCLE gene expression dataset, we impose sparsity constraints on the *H* matrix. We use this matrix to define cell states via clustering, making it essential to find a good balance between local and global transcriptional structures. In the second panel of Fig. 2A and 2B, we observe that the standard *NMF* algorithm tends to generate representations favoring global patterns. For the faces example, this means the image components matrix (*W*_*n,k=80*_) displays patterns consistent with complete faces. In the CCLE example, the patterns correspond to sample component matrix (*H*_*k=16,m*_) over-expression patterns that involve a significant fraction of the cancer samples. In contrast, imposing strong sparsity constraints on the *W* (top) or *H* (bottom) matrices shifts the *vsNMF* solution to favor highly localized features. Under these constraints, the feature components matrix (*W)* contains more isolated image “parts”, such as individual mouths, noses, and eyes, in the faces dataset. Meanwhile, the sample component matrix (*H*) in the CCLE dataset reveals much more localized over-expression patterns, which may involve only samples from a specific subtype. In the third panel of Fig. 2A and 2B, we apply a power-law sparsity function, which yields a combination of local and global patterns with an emphasis on the sparser, more localized ones. For the faces dataset, this results in some entries within the image component matrix resembling entire faces, while others look like isolated facial parts. This transition is visible in the way the initial entries of the faces in the *W* matrix are mostly dark but become progressively whiter further along the figure. A similar arrangement appears in the CCLE example, where some gene expression patterns are global, much like in standard *NMF*, while others are highly localized. The balance of local versus global (dense versus sparse) components can be tuned to better suit the goals of the dimensionality-reduced model representation. We believe this ability to control the mixture is crucial for making the dimensionality-reduced projection more representative of the hierarchical structure inherent in genomic datasets, thereby making it easier to interpret biologically.

**Figure 2.**
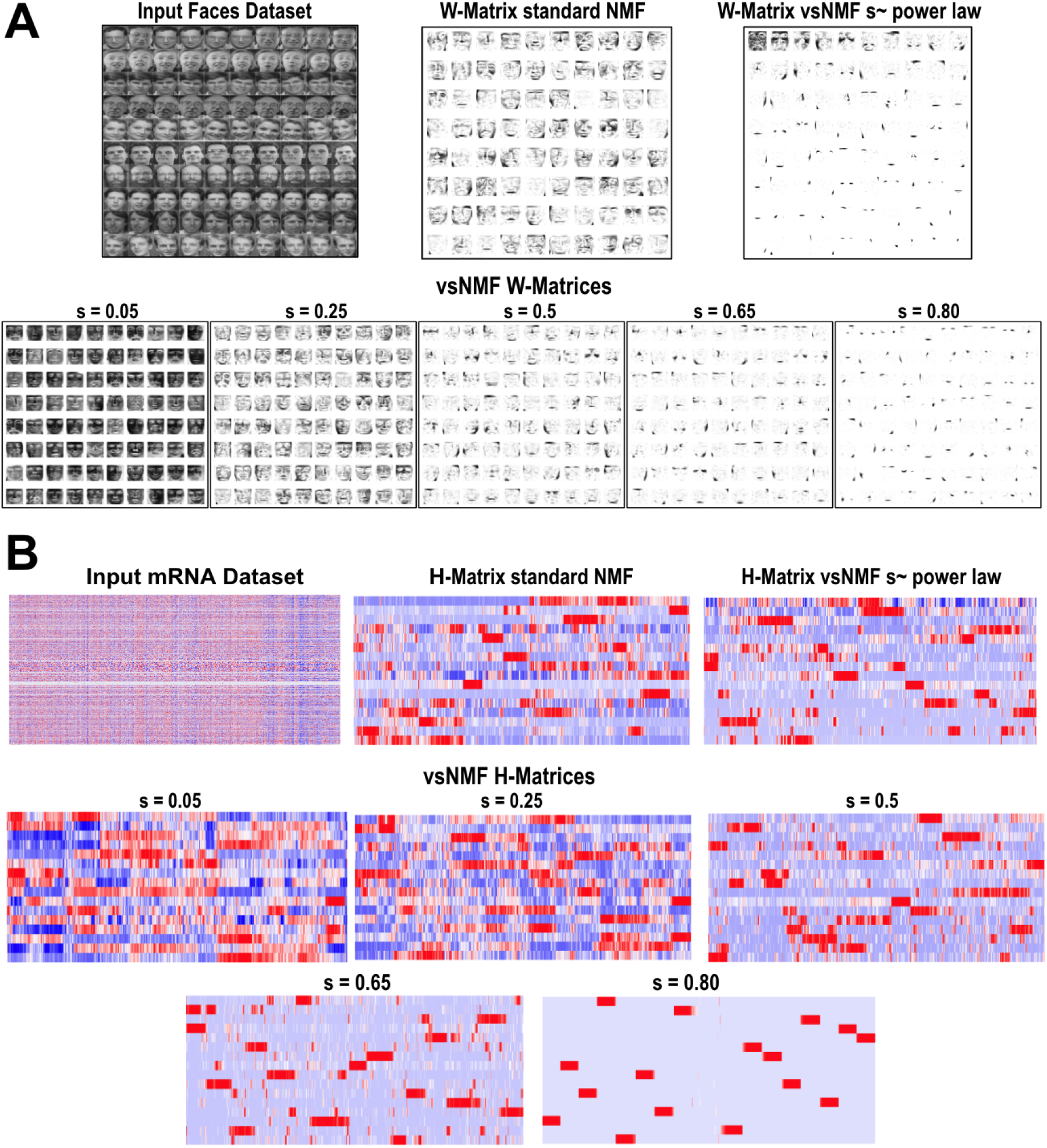
Standard *NMF* and *vsNMF* Factorizations for (A) ORL Images and (B) *CCLE* mRNA dataset.

The additional sparsity-enforcing projective step in *vsNMF* increases the computational cost of each iteration. Consequently, convergence is slower, as illustrated in Fig. 3A, resulting in total execution times that can be up to an order of magnitude longer (Fig 3B). Despite this increase, the longer runtime remains acceptable for most use cases and does not diminish the overall practical utility of the approach.

**Figure 3.**
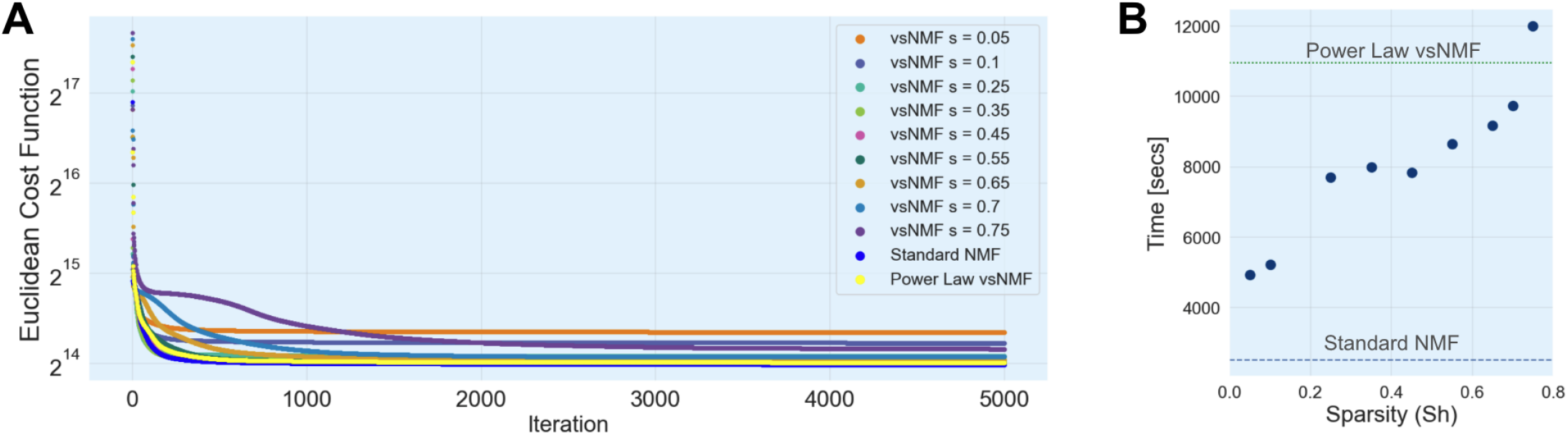
A) convergence over time of standard NMF and vsNMF with different sparsities, and B) the corresponding total computer execution times.

### Projecting new samples onto an existing vsNMF model

An independent dataset *B* can be projected onto the *vsNMF* model space to obtain a projected version of the dataset in the space of the transcriptional

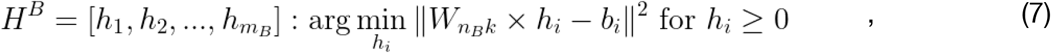

signatures in the same way as in standard *NMF*. This is accomplished using a *non-negative least squares solver* [49] where we solve a linear problem for each column of *B*,where *n*_*B*_ is the set of genes in common between *B* and the original *reference* dataset *A* where the transcriptional signatures were generated. This projection can also be implemented by using the Moore-Penrose pseudo-inverse of the *W* matrix [20],

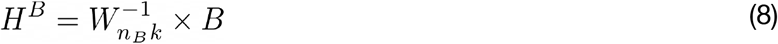

and truncating the negative elements to zero. Once projected, the sample profiles (*H*^*B*^) from dataset *B* can be compared with each other, or with other datasets’ projected samples.

### Examples of *vsNMF* applications and comparisons with standard *NMF*

In this section we present two examples of the use of *vsNMF* in functional cancer genomics problems. These examples will illustrate the benefits of having a much greater control of the desired sparsity imposed on the *NMF* solution. There is a third example that we will present after the introduction of the vsNTF algorithm.

### Example 1. Defining *vsNMF* transcriptional signatures that align with single genes mRNA profiles

In functional genomics, *NMF* is frequently used to create low-dimensional representations or embeddings of transcriptional data, where the rows of the *H* matrix act as transcriptional signatures that summarize the dataset. These signature values can be used to represent, compare, and cluster samples within this lower-dimensional space. In this context, it is often desirable for the rows of the *H* matrix to closely match the actual expression profiles of individual genes. This alignment between rows of the *H* matrix and the input profiles is particularly useful when annotating and characterizing these signatures by comparing them against the profiles of genes or known gene sets, for example, using single-sample GSEA [50]. Because standard *NMF* does not necessarily favor this type of solution, one might ask whether adjusting the sparsity functions can indirectly control this behavior. Specifically, one may envision using the sparsity function of *vsNMF* to control the *H* matrix to become more convex-like without enforcing the strict mathematical constraints required by feature-convex *NMF* [51]. As we will show below, this is indeed possible by using *vsNMF* and specifying an appropriate sparsity function (*s*_*H*_(*k*)) that favors a more global structure for the resulting transcriptional signatures.

In this example we decompose solid tumor cell lines mRNA profiles from the Cancer Cell Line Encyclopedia (*CCLE* [47,48]) compendium in order to generate transcriptional signatures that summarize the overall patterns of gene expression. To prepare the dataset we replaced the mRNA values with their column-ranks to make each sample have the same dynamic range and reduce the effect of outliers. We also filtered genes to define a subset of oncogenic-related genes and minimize the effect of tissue markers in the signature generation and clustering steps, and at the same time maximize the detection of common oncogenic pathways across cancers. We implement this strategy by using a subset of 3,958 oncogenic-related genes that were identified from oncogenic activation isogenic assays [13] for 14 major oncogenes (*KRAS, ERBB2, ZEB1, AXL, MYC, CTNNB1, BRAF, ETV1, PIK3CA, EGFR, ESR1, AR, PGR*, and *NFE2L2*), that define a suitable universe of common pan-cancer oncogenic activation. Fig. 4 shows the overall methodology.

**Figure 4.**
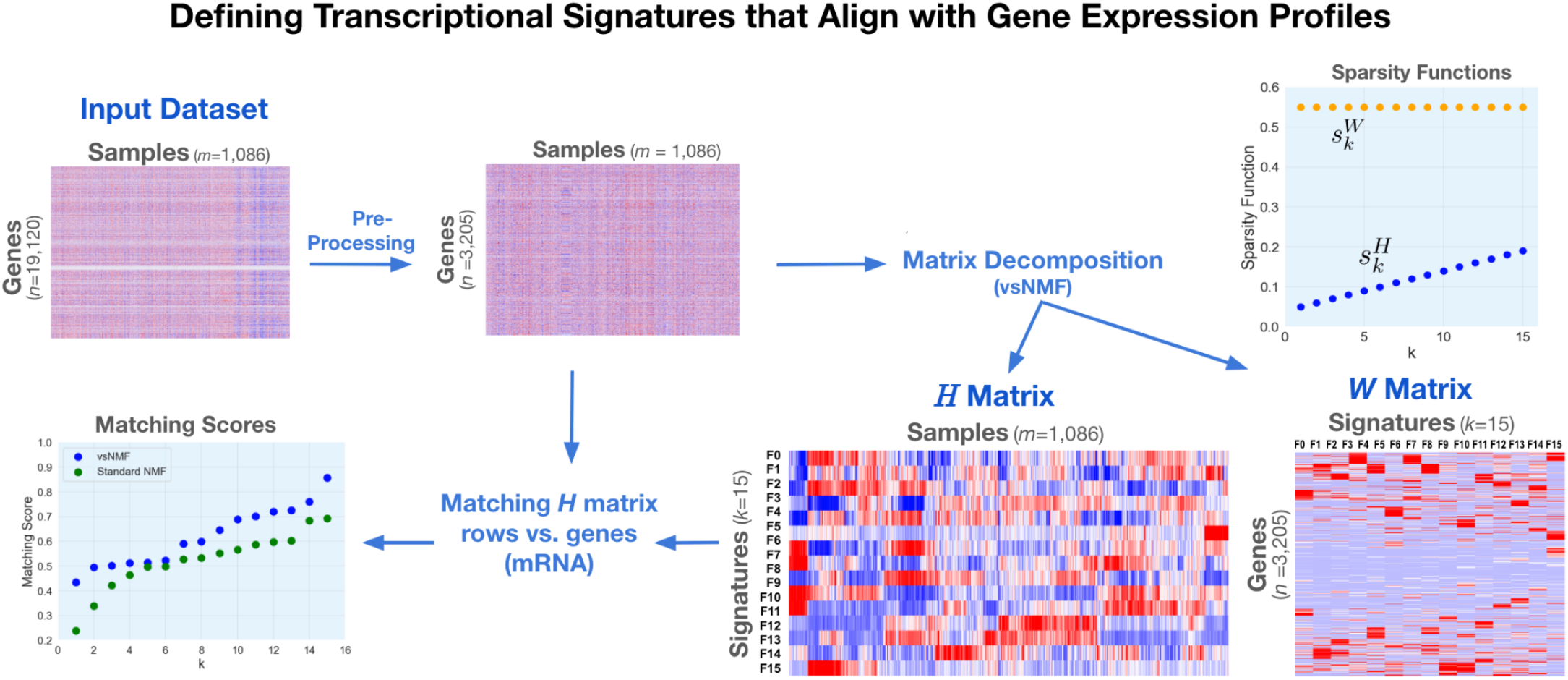
Methodology to generate transcriptional signatures that summarize the overall patterns of gene expression in pan-cancer cell lines using *vsNMF* and a specific *H*-matrix low sparsity function.

A preliminary analysis indicated that lower sparsity on the *H* matrix, and an intermediate sparsity on the *W* matrix, will favor better matching between the signatures (rows of *H*) and gene expression profiles from the input dataset. Based on this observation we ran the vsNMF algorithm and specified lower sparsities for *H* following a linear function from 0.05 up to 0.20, and a fixed sparsity of 0.55 for *W* (Fig. 4). Using these sparsity functions we ran *vsNMF* for 1,000 steps using 15 components (signatures). The resulting *W* and *H* matrices are shown in Fig. 4.

Once the *vsNMF* run was completed we matched each of the *H* matrix rows vs. the mRNA profiles of the input dataset (3,958 genes) and averaged the matching scores of the top twenty genes per signature component. The matching score used in this analysis is the *Information Coefficient* (IC) [52–54] which is a rescaled version of the mutual information that has values between -1 and 1 like a correlation coefficient, but can measure statistical dependency between variables associated with for linear and non-linear relationships. The results are shown in Fig. 5 where they are compared with the corresponding scores for the matching of the *H* matrix rows of a standard *NMF* run using the same input dataset and the same number of steps (1,000) and components (15). As can be seen in the figure, the use of specific sparsity constraints, and in particular imposing low sparsity constraints on the *H* matrix, allowed the *vsNMF* algorithm to significantly improve the matching scores of all the components in comparison with standard *NMF*. A more systematic exploration of sparsity functions may be able to improve the results further.

**Figure 5.**
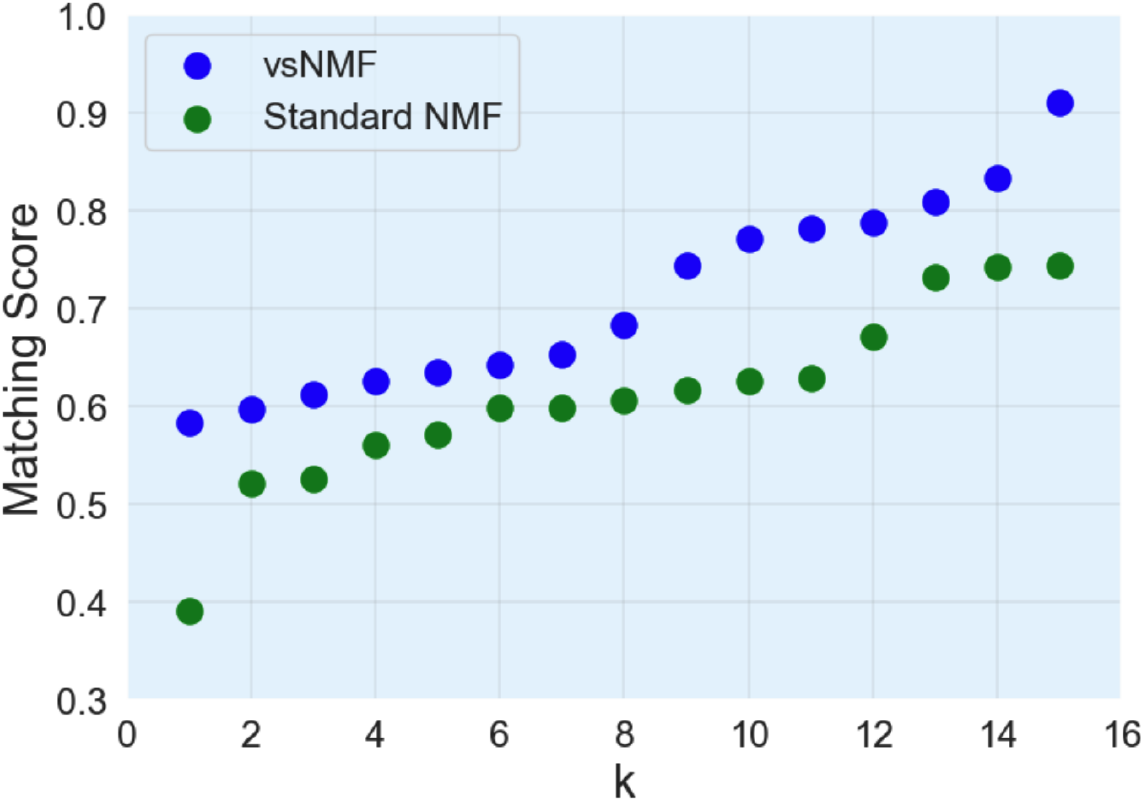
Matching scores (IC) from matching each of the *H* matrix rows vs. the mRNA profiles of the input dataset and averaged the matching scores of the top twenty genes per signature component for *vsNMF* and standard *NMF*.

### Example 2. Clustering samples to define improved groups or putative cell states

In many cancer genomic applications the samples of an input dataset are clustered in the space of *NMF*-defined signatures for the purpose of defining sample groups, subtypes or putative states [13,55]. Under this approach it is desirable to generate clusters that represent good clustering solutions in the sense of lowering distorsion, increase reproducibility and robustness, and improved cluster compactness and separation. As the standard *NMF* algorithm does not necessarily optimize the *H* matrix solution to achieve this goal, we will use the sparsity functions in *vsNMF* to maximize the quality of the clustering solution and use two clustering metrics to evaluate the results: the *cophenetic coefficient* [56] and the *silhouette inde*x [57]. The cophenetic correlation coefficient measures how faithfully a clustering solution preserves the original pairwise distances between the data points. It works by calculating the correlation between the original distance between two data points, e.g., their Euclidean distance, and the “dendrogram distance,” i.e. the height of the tree at which those two points are first merged into the same cluster. It ranges from 0 to 1, and values close to 1 indicate that the clustering solution is a highly accurate reflection of the true data structure. Lower values suggest that the clustering algorithm is forcing the data into a structure that does not naturally fit the data. The silhouette index ranges from -1 to 1, and measures cluster cohesion versus cluster separation, i.e. how similar a data point is to its own cluster compared to other clusters. For every single data point, it calculates the average distance between the point and all other points in its own cluster (cohesion), and the average distance between the point and all points in the nearest neighboring cluster (separation). Values near 1 indicate that a data point is far away from neighboring clusters and very close to members of its own cluster. Values near 0 indicate that a data point is on or very close to the decision boundary between two neighboring clusters, and values near -1 show that a data point is closer to a neighboring cluster than to its own cluster and it is being possibly misclassified. The final measure is the average silhouette score across all data points and provides a global measure of how distinct and well-separated the clusters are and a high average silhouette score indicates a strong, well-defined clustering solution.

For this example we will use the same input dataset as in Example 1, representing the transcriptional profiles of solid tumor pan-cancer cell lines, and use *vsNMF* to generate transcriptional signatures and cluster them using hierarchical clustering with Ward linkage. Because hierarchical clustering requires a distance metric rather than a similarity metric, we use one minus the *Information Coefficient* (IC) [52–54] for each pair of samples (columns) of the *H* matrix to generate a distance matrix so that high mutual information becomes a small distance. In this way the distances are bounded by [0, 2] where 0 represents no distance, as e.g. when comparing an *H* Matrix column with itself, and 2 represents maximum distance, as when an *H* matrix column is completely anti-correlated with respect to another one. Using this distance matrix, the clustering algorithm performs (bottom-up) hierarchical clustering using Ward’s minimum variance method [58] and defines 10 groups. Ward’s method takes an analysis-of-variance approach and instead of just looking at the distance between the edges of clusters, it evaluates the “center of mass.” In this way at every step, it merges the two clusters that will result in the smallest possible increase in the total within-cluster variance, i.e. the error sum of squares. This mathematical objective forces the algorithm to build tight, cohesive, and well-balanced groups, which is especially useful when trying to identify distinct subtypes or robust sample groups. As we don’t know in advance what would be the best settings for the sparsity function we will explore using different values for constant *s*_*H*_ functions in the range [0.05, 0.80] and keeping *s*_*W*_ unconstrained. The methodology is shown in Fig. 6. In Fig. 7A we can see that higher values of the *H*-matrix spasticity function, above 0.6, results in increased silhouette indices and increased quality of the clustering solutions. The standard *NMF* silhouette index is 0.13 for this dataset (Fig. 7B), and when we use vsNMF with a constant *H*-matrix sparsity of 0.65, we can increase the silhouette index to 0.251 (Fig. 7C), which is about twice the standard *NMF* value. This is a significant improvement in the quality of the clustering solution. Consistent with these results, the cophenetic coefficient also increases for constant *H*-matrix sparsity above 0.6.

**Figure 6.**
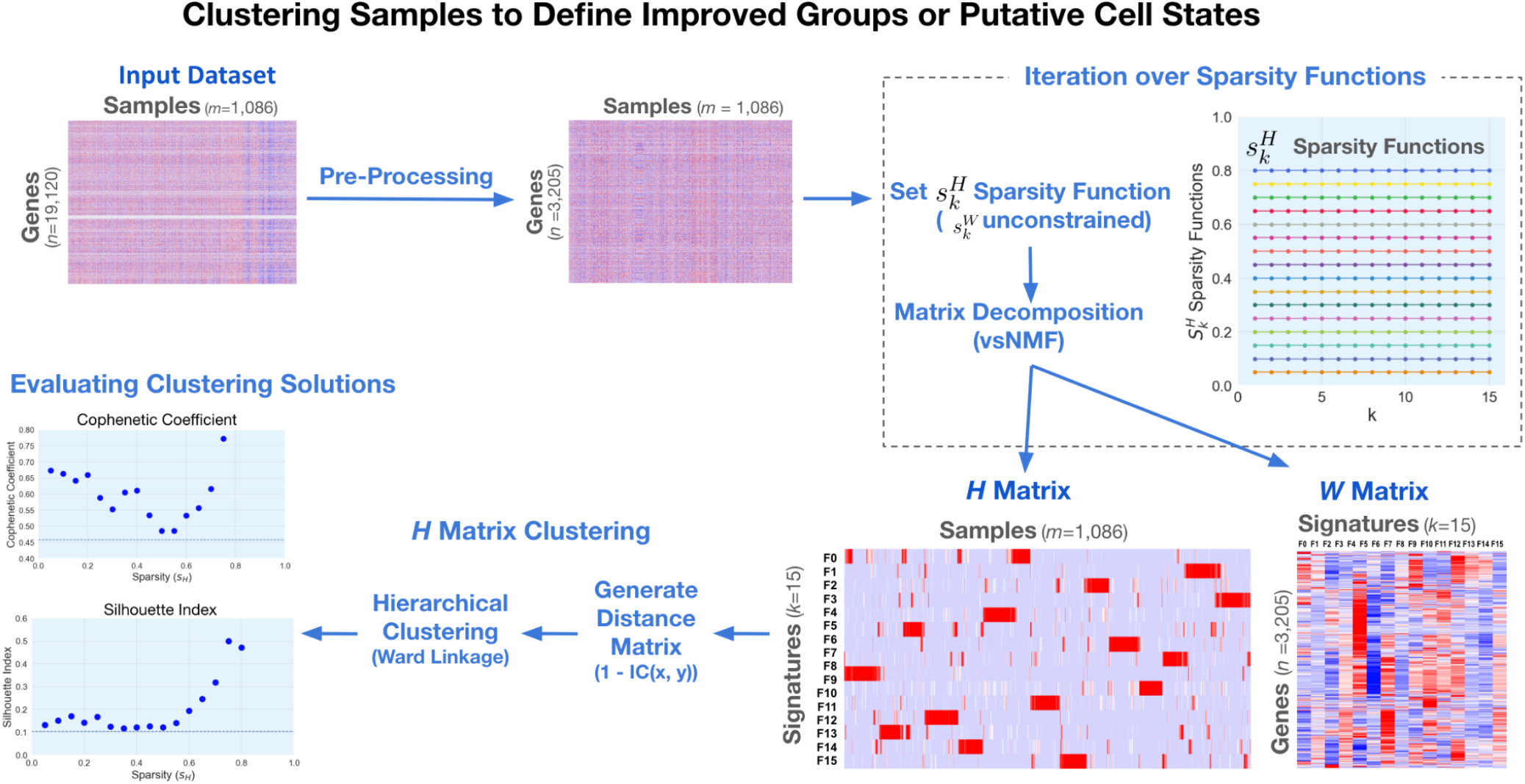
Methodology to cluster *vsNMF*-projected samples to define improved clustering solutions.

**Figure 7.**
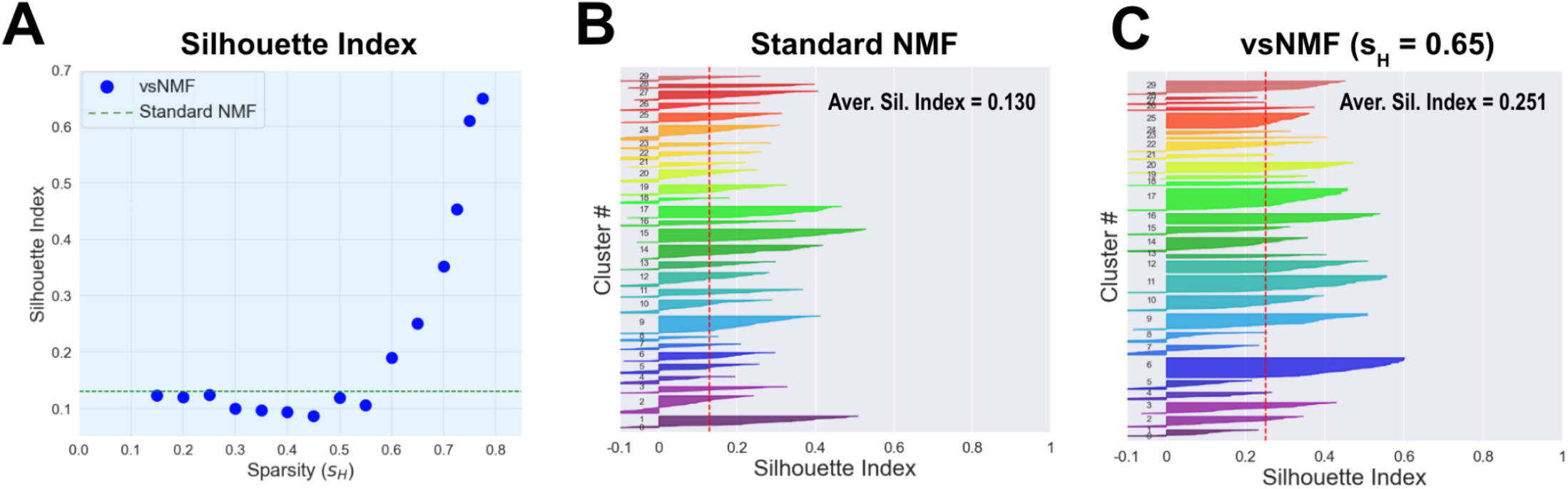
A) Silhouette index as a function of *vsNMF H*-matrix spasticity function values, B) standard *NMF* silhouette index, and C) silhouette index for *vsNMF* with a constant *H*-matrix sparsity of 0.65.

As in the previous example, a more systematic analysis of the sparsity function profiles could result in further improvements of the clustering solution. In standard *NMF*, dense *H* matrices can blur the boundaries between distinct biological groups, leading to overlapping and ambiguous clusters. By enforcing higher sparsity on the sample embeddings (*H* matrix), *vsNMF* effectively filters out background noise and shared, non-specific signals, forcing the model to represent samples using only their most defining, “localized” features. This targeted dimensionality reduction inherently produces tighter, more cohesive, and better-separated clusters. Ultimately, the flexibility to tune these sparsity constraints makes *vsNMF* a highly versatile tool for discovering robust, biologically meaningful subtypes in complex, high-dimensional genomic data, overcoming the rigid, often overly global representations produced by standard NMF.

### The *variable-sparsity Non-Negative Tensor Factorization* algorithm (vsNTF)

Tensors are a higher dimensional (three or greater) generalization of matrices, just as matrices are a two-dimensional generalization of vectors. The higher dimensionality of tensors means that they inherently preserve complex multi-factorial interactions that are often lost when representing the data as a 2-dimensional matrix. Like matrices, tensors can also be decomposed to reveal latent factors and to perform pattern discovery and define groups of genes, samples and phenotypes with greater similarity. The primary methods for tensor factorization are Canonical Polyadic Decomposition (*CPD*) and Tucker Decomposition [59]. *CPD* breaks an input tensor into a sum of tensor products of vectors representing each of the input tensor’s dimensions [60]. The vectors in these tensor products can be organized as columns of factor matrices [61]. Wang et al. applied *CPD* to scRNAseq data from different tissue types and showed superior cluster separation compared to the classic 2-dimensional *PCA* analysis applied to a matricized form of the tensor [62]. In Tucker tensor decomposition, an *N*-dimensional input tensor is factorized into a smaller, dense core tensor and a set of factor matrices, specifically, one factor matrix for each dimension of the tensor. The factor matrices are analogous to the *W* and *H* matrices in *NMF* since they describe how much each actual feature represents each meta-factor (see Fig. 8A). The core tensor acts as a compressed version of the original input tensor and captures the interactions between the different components in the factor matrices. The factor matrices extract the latent features or patterns along each specific dimension, while the core tensor shows how strongly these latent features interact with one another. Tucker tensors are often used for data compression but can also be used for data analysis, and *CPD* can be considered a special case of Tucker factorization where the core tensor is simply an identity tensor [61]. Normally, Tucker Decomposition side matrices are hard to interpret since they are non-unique; however, it is possible to impose non-negativity constraints on Tucker Decomposition to make these factors interpretable for use in genomic data analysis [61].

**Figure 8.**
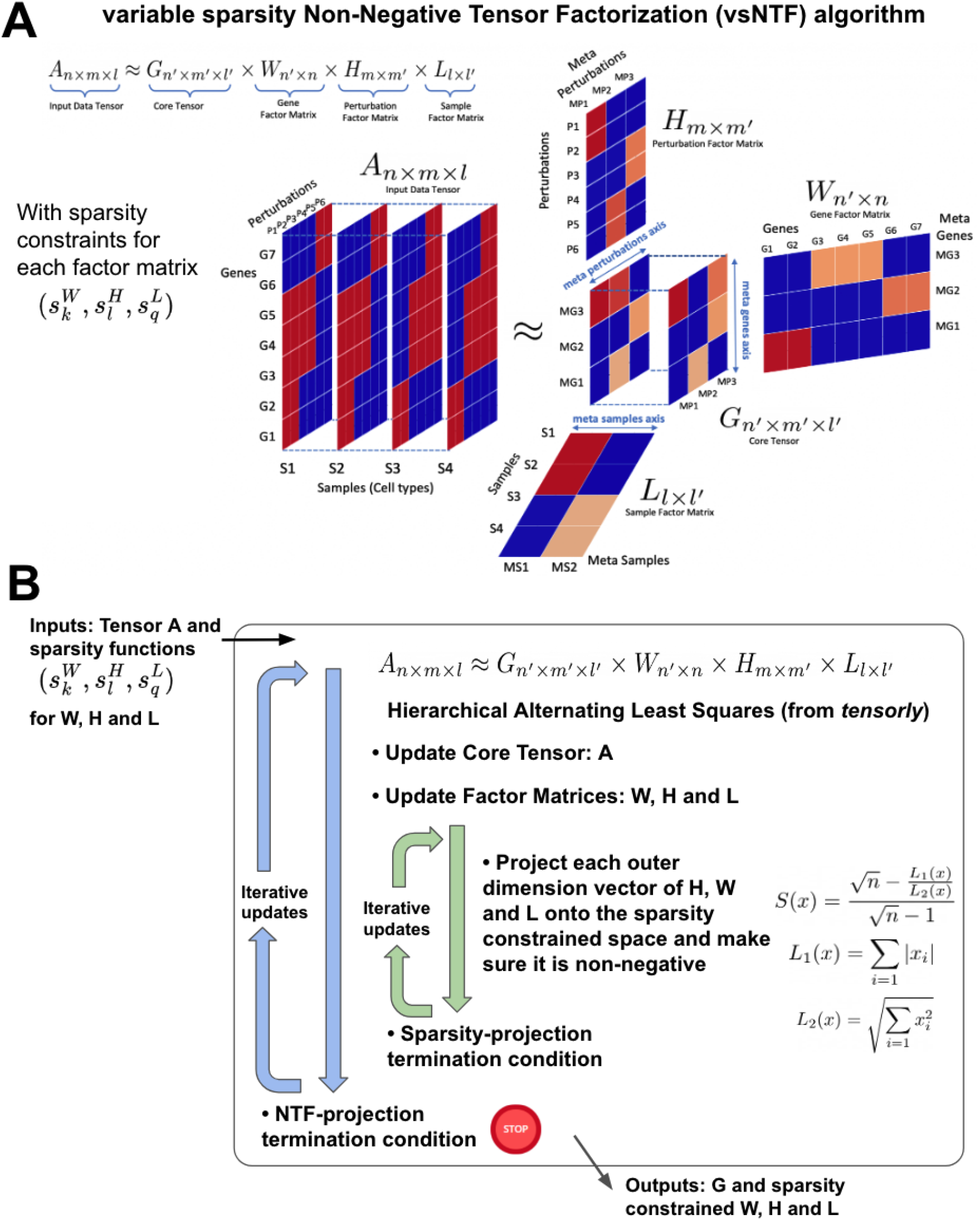
A) Non-Negative Tucker Tensor decomposition, and B) top-level description of the *vsNTF* decomposition algorithm.

Non-negative Tucker Factorization *(NTF*), sometimes called Non-negative Tucker Decomposition, is simply Tucker decomposition with non-negativity constraints imposed on the core tensor and all the factor matrices involved. This non-negativity constraint allows the factors encoded in the factor matrices to be more interpretable. *NTF* is a direct higher order generalization of *NMF*. In fact, *NMF* can be considered a special case of *NTF* where the core tensor is an identity matrix, a second order tensor, and *W* and *H* are the two factor matrices [63]. The multiplicative update algorithm from *NMF* can also be used to iteratively solve for the core tensors and factor matrices in *NTF*. Sparsity constraints have also been proposed as a way to gain more local scale components and improve interpretability [63], however, most sparsity constraints are simply based on the *L1* norm, which enforces sparsity but does not take into account global features the way the *L1/ L2* ratio encourages scale-invariant solutions [64]. *NTF’*s re-use of multiplicative update also allows us to impose *L1/L2* ratio based sparsity constraints in the same way we imposed on the different components of *vsNMF*’s *W* and *H* matrices. To implement our variable sparsity Non-Negative Tensor Decomposition (*vsNTF*), we modified the Non-negative Tucker decomposition function from the *Tensorly* package [65], specifically to impose the *L1/L2* norm sparsity-based projection after every multiplicative *NTF* update iteration (Fig. 8B). The sparsity constraints can be uniquely defined for every inner factor of the side matrices as was imposed on the *W*-columns and *H*-rows in *vsNMF*.

### Example 3: Applying *vsNTF* to analyze COVID data and detect cytokine storms

We applied *vsNTF* to a dataset of COVID patients published by Ren et al. 2021 [66]. This dataset contains single cell RNAseq profiles corresponding to 27,943 genes from 196 patients hospitalized for COVID-19 infection in China. There were a total of 1,462,702 cells from 64 cell types, of which 4 were lung epithelial cells and 60 were immune cells, whose labels were assigned based on clustering the data in 2D matrix form and visualizing expression patterns of known histological marker genes [66]. A graphical summary of the data is shown in Fig. 9.

**Figure 9.**
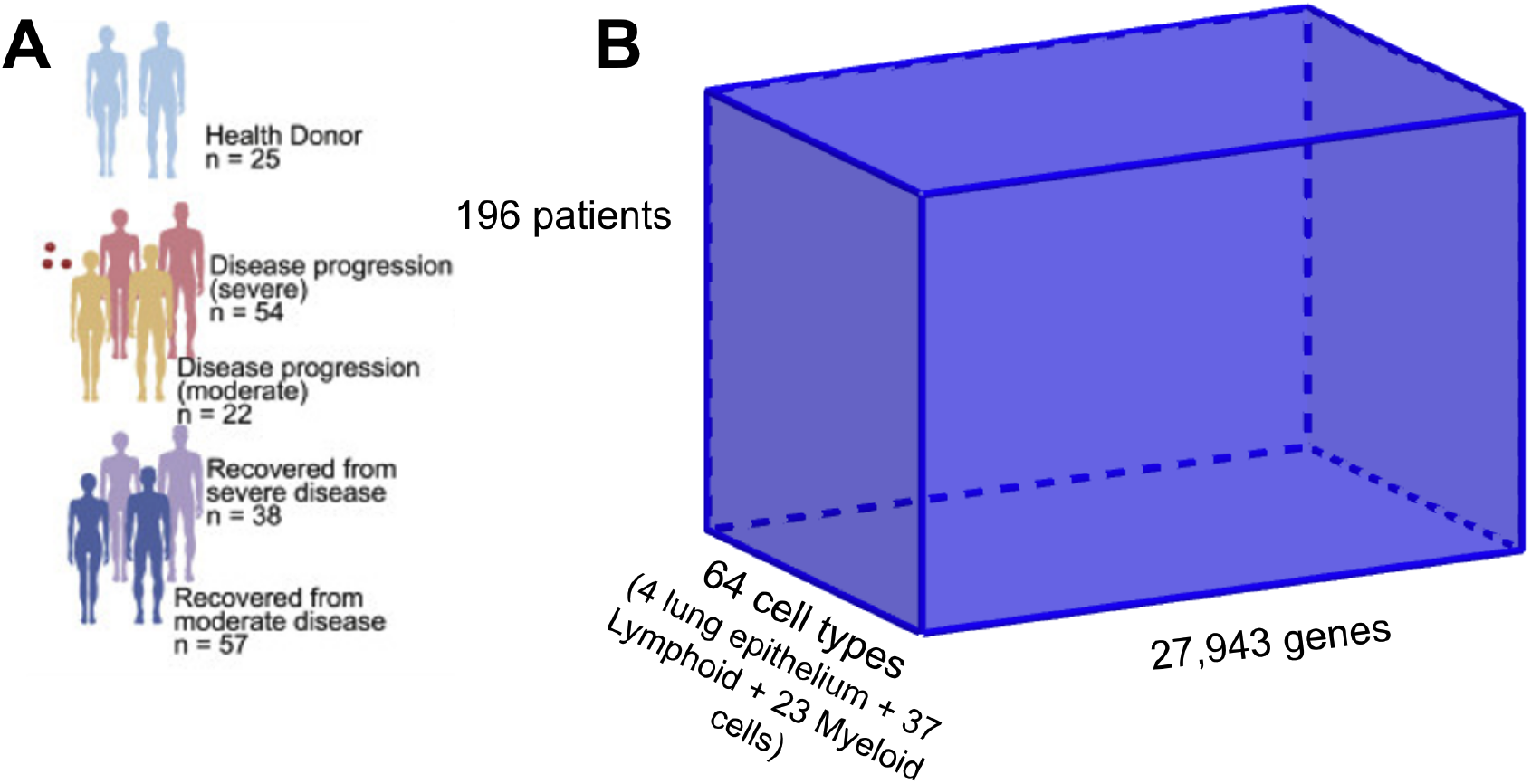
Graphical summary of the COVID-19 single cell dataset (A) and the resulting scRNAseq tensor (B).

Cytokine storms occur when immune cells release high concentrations of pro-inflammatory cytokines, often triggered by the presence of a pathogen such as a virus or a bacteria. While inflammation is part of the normal immune system’s defensive process, these cytokine storms often result in excessive inflammation and tissue damage. Cytokine storms are thought to be the cause of COVID-19’s symptoms and the reason for severe clinical symptoms in certain patients. Ren et al’s analysis of their scRNAseq COVID data set showed Megakaryocytes, certain types of monocytes (Mono_c1-CD14-CCL3, Mono_c2-CD14-HLA-DPB1, and Mono_c3-CD14-VCAN), and some subtypes of T-cells (T_CD4_c08-GZMK-FOShigh, T_CD8_c06-TNF, and T_CD8_c09-SLC4A10) as the main source of cytokines in some severe COVID cases. While the initial analysis [66], was performed on a 2D matrix, it is possible to reconfigure this data as a 3D tensor with dimensions: patient x gene x cell type. scRNAseq data is typically very sparse, so we filtered out genes where over 70% of cells were zero-valued, leaving 11,273/27,943 genes. Since each patient has a different number of cells of each type, we took the average expression of all genes in all cells of the same type for each patient. The final dimensions of the tensor were: 196 patients x 11,273 genes x 64 cell types (Fig. 9B). This tensor was decomposed using *vsNTF* and selecting internal factor dimensions as follows: 8 meta-patients (patient groups) x 12 meta-genes x 10 cell type groups. The variable sparsity constraint, again using a power law based function [67,68], was imposed only on the meta-genes since it corresponds to the dimension with the highest number of features. In order to make a fair comparison, standard (non-sparsity constrained) *NTF*, using the same internal factor dimensions, was also used on the tensor to evaluate the effects of the variable sparsity constraints.

To identify which metagene was responsible for the cytokine storms, we examined the relative amplitudes of cytokines shown in Fig. 6E of Ren et. al 2021: *PDGFA, TGFB1, TNFSF4, PF4V1, PPBP, TNF, CCL4, CCL4L2, IL1RN, CXCL8, CCL3L1, IL1B, CXCL2, CCL3, IL6, SCYL3, IL18, HBEGF, TNFSF10, TNFSF13, FTH1, TNFSF13B, CXCL16, CD40LG, LTB, TNFSF8, FGF23, IGF1, TXLNA, SCYL1, IL16, VEGFB, MIF, CCL5, IL32, CD70, IFNG, XCL2*, and *XCL1* of the metagene factor matrix (Fig. 10B). MG2 and MG3 both have relatively high amplitudes in the genes corresponding to hyper-inflammatory myeloid cells: Mono_c1-CD14-CCL3, Mono_c2-CD14-HLA-DPB1, and Mono_c3-CD14-VCAN in Fig. 6E of *Ren et. al 2021* [66] (Fig. 10B).

**Figure 10.**
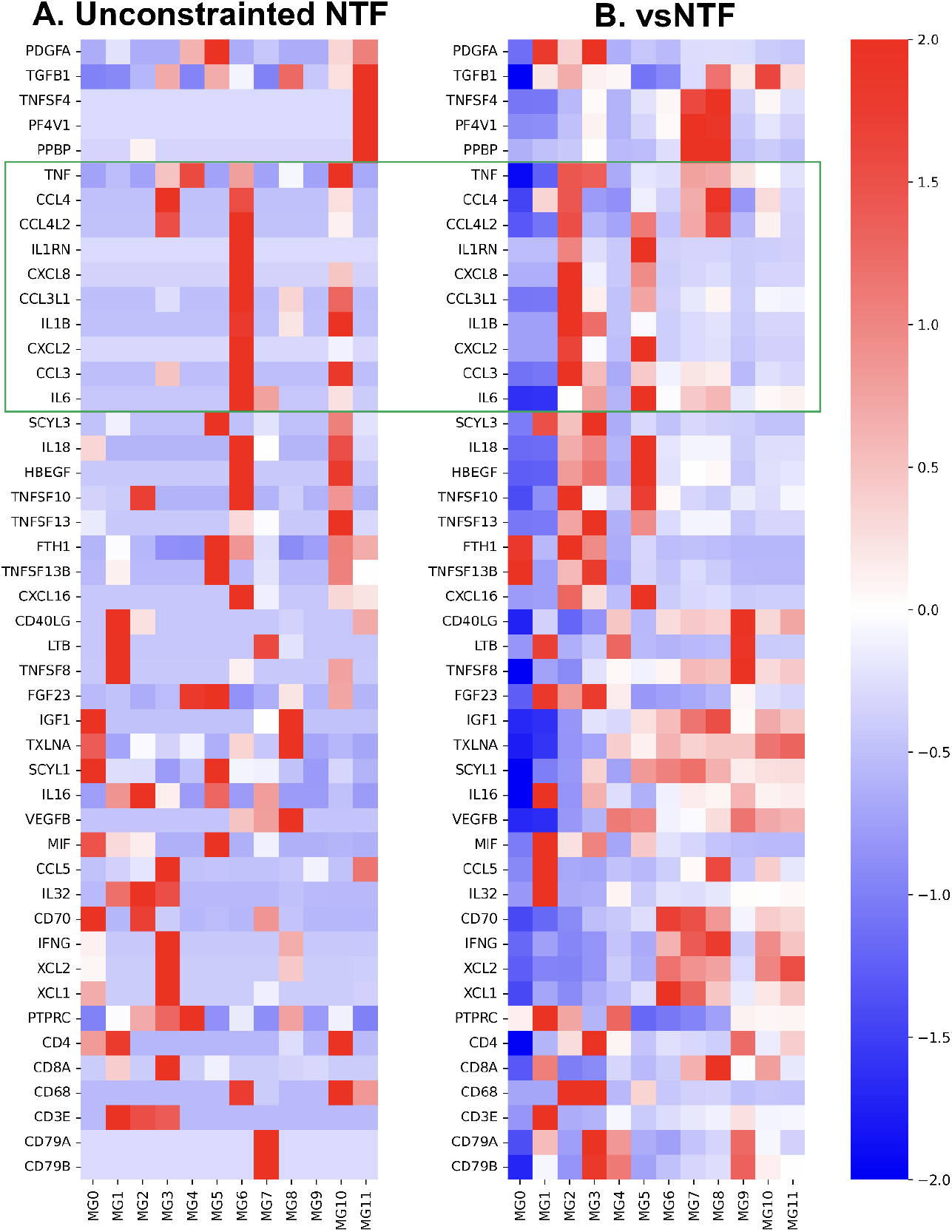
Side matrix slices of the metagene factor matrix showing relative amplitudes of cytokines in Figure 6E of the *Ren et al. 2021* dataset and some histological cell type marker genes. The green box indicates genes associated with hyper-inflammatory myeloid cells in Ren et al.’s analysis. A). The side matrix in unconstrained *NTF*. B) The side matrix in *vsNTF* shows consistently higher MG2 and MG3 amplitudes in genes associated with hyper-inflammatory myeloid cells. The color scale at the far right represents the Z-scores.

Next, we examine the cross-dimensional relationship between certain meta-factors and actual features. Cross-dimensional relationships between meta-factors from one dimension and actual features from another dimension can be explored by multiplying the core tensor, with the appropriate factor matrix, then unfolding the tensor. First, we examine the relationship between metagenes and cell types. Fig. 11B shows that MG2 and 3 are highly enriched within the hyper-inflammatory myeloid cells, confirming the observation in the metagene side matrix (Fig. 11B). We analyzed the metagene to patient relationship by multiplying the core tensor with the metapatient factor matrix and unfolding the tensor product to determine the clinical effects of each Metagene. Since the data is *in-vivo*, with many confounding variables, and we are limited by the small subset of clinical variables that the authors made available to the public, it is not immediately clear what the exact clinical significance of MG2 and MG3, or any of the other metagenes are. However, we observe that MG1 is significantly associated (p < 0.05) with less severe disease and with the healthy controls using the Mann-Whitney test. MG1 appears to be associated with lymphoid cells, which all show a moderately high expression of its genes. This corresponds to the lymphopenia, or lack of T-cells, observed in severe COVID-19 cases [69], though this means that the metagene might simply be a lymphoid signature, not representative of any precise biological activity.

**Figure 11.**
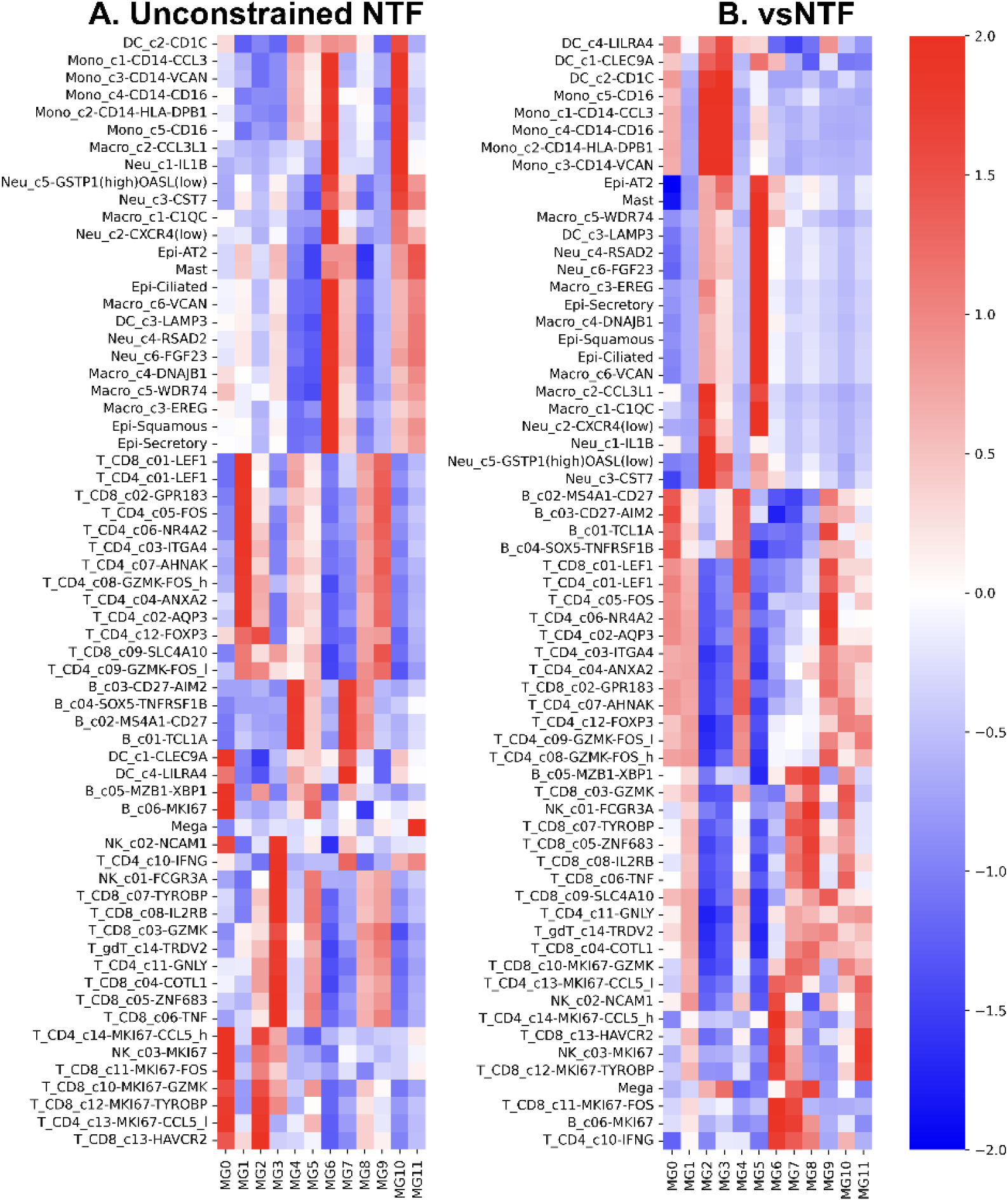
Relationship between metagenes and cell types in A. unconstrained *NTF* and B. *vsNTF*. The color scale at the far right represents the Z-scores.

The results of performing the same decomposition but using standard *NTF* without sparsity constraints show that it is also able to discover a metagene, MG10, associated with Cytokine storms. However, it is less specific than MG2 and MG3 in the *vsNTF* components since it has a relatively high amplitude in more cell types. Genes associated with hyper-inflammatory myeloid cells also do not show consistently high values in MG10 of unconstrained *NTF* as they do in MG2 and 3 of the *vsNTF* metagene matrix. MG6 in the unconstrained *NTF* shows consistently high expression in those genes, but the hyper-inflammatory myeloid cell types show a paradoxically low amplitude in MG6 when using unconstrained *NTF*. In addition, MG6 tends to have high values in most myeloid cell types, indicating it might simply be a myeloid cell type signature in a similar way MG1 in the *vsNTF* components appears to be a lymphoid signature. This means that myeloid cells’, mainly associated with non-specific innate immunity, and thus a larger variety of cellular activities due to the multiplicity of different pathogens, expression patterns are more local scale than the lymphoid expression patterns, which are optimized for responding to specific antigens [142]. Improved tuning of the variable sparsity constraints could allow *vsNTF* to recover more meaningful metagenes in both myeloid and lymphoid cell types.

## Discussion and conclusions

This paper introduces *vsNMF*, a variable-sparsity extension of *NMF*, and non-negative Tensor factorization (*vsNTF*), that allow users to prescribe a separate sparsity level for individual components, the columns of *W* and the rows of *H*, or factors along tensor modes. Unlike methods that add sparsity penalties to the *NMF*’s objective function, these algorithms directly enforce sparsity through a projection step after each factor update while retaining reconstruction-error minimization as the underlying objective. In this way, *vsNMF* strictly preserves the fundamental reproduction error minimization objective of standard *NMF*, but its primary innovation lies in introducing explicit, fine-grained control over the sparseness of the factorization. This architectural refinement is highly significant because it allows researchers to directly modulate the algorithm’s inherent representational duality without distorting the underlying objective function. Rather than being locked into a single mode of interpretation, *vsNMF* enables the user to smoothly interpolate the solution along a geometric continuum. At the level of individual components, the algorithm can be tuned to operate anywhere from an “extreme-ray” regime, yielding a generative, parts-based encoding of overlapping features, to a “centroid” regime, which behaves as a partitional, soft-clustering model. Consequently, *vsNMF* provides unprecedented flexibility to tailor the specific mixture of global and local structures to the unique demands of diverse genomics use cases. Assigning different sparsity levels, e.g. through a power-law distribution of sparsities, can produce a multiscale representation containing both broad and specific biologically-relevant patterns. The community currently relies heavily on dimensionality reduction techniques like *UMAP* or t-SNE for visualization, which often distort global structures, or standard *PCA*, which can lack biological specificity. In the application examples, we show how one can achieve better signature alignment against gene expression profiles than standard *NMF*, resulting in “convex-like” behavior without strict convexity constraints. We also show how sparsity can be tuned to optimize clustering quality, e.g. the silhouette index, for defining groups of cancer samples as putative states. The *vsNTF* example shows the decomposition of COVID19 data where specific isolated metagenes captured the hyper-inflammatory myeloid cells driving cytokine storms, whereas unconstrained *NTF* produced muddled or paradoxical signatures. A specific lymphoid-associated metagene was linked to milder disease.

The availability of the *vsNMF* and *vsTNF* algorithms provides the cancer genomics and bioinformatics community with several benefits. The algorithms offer much greater flexibility in decomposing datasets and controlling global and local structures of the solutions. The flexibility provided by variable sparsity to maximize or minimize any measurable property of phenotype could find wide applicability in many problems. The *vsNMF* and *vsNTF* learned representations can serve as reusable reference spaces or embeddings where new datasets can be projected into an existing *vsNMF* or *vsNTF* model using non-negative least squares or a truncated pseudoinverse. This provides a method to project samples from different experiments, including single-cell data, to a suitable common projection space where the chosen sparsities maximize the biological relevance of the learned representation. By providing a “tunable” dimensionality reduction technique, the algorithm could provide dimensionality-reduced representation models that are not only useful for visualization but are better at representing the underlying biology and allow to model many datasets in a flexible, unified and interpretable way.

## Acknowledgments

The work presented here was supported in part by NIH grants: R01HG009285, R01CA154480, R01CA121941, R01CA247551, R01DE026870, U24CA220341, U24CA248457, R01CA226803, U01CA217885, P30CA023100, R01CA109467, R01DE035393, R01CA292590, U24CA295532, P01CA281819, U01CA268052, U01CA278470, R01NS122835, R01DE026644, R01CA257505, U54CA209891, U01DE028227, a Cancer Target Discovery and Development (CTD2) grant (U01CA176058) and a State of California’s Initiative to Advance Precision Medicine (CIAPM) award (OPR18112).

## Software

The python programs that implement the computation of the *vsNMF* and the *vsNTF*, and a Jupyter notebook with examples, are available at https://github.com/pablotamayo/vsNMF. The *vsNMF* code is based on an original MATLAB program by Patrik Hoyer (github.com/aludnam/MATLAB/blob/master/nmfpack/) and described in [37]. The *vsNTF* code is a modification of the original tensorly package [65](github.com/tensorly/tensorly)

